# Using predictive specificity to determine when gene set analysis is biologically meaningful

**DOI:** 10.1101/080127

**Authors:** Sara Ballouz, Paul Pavlidis, Jesse Gillis

## Abstract

Gene set analysis, which translates gene lists into enriched functions, is among the most common bioinformatic methods. Yet few would advocate taking the results at face value. Not only is there no agreement on the algorithms themselves, there is no agreement on how to benchmark them. In this paper, we evaluate the robustness and uniqueness of enrichment results as a means of assessing methods even where correctness is unknown. We show that heavily annotated (“multifunctional”) genes are likely to appear in genomics study results and drive the generation of biologically non-specific enrichment results as well as highly fragile significances. By providing a means of determining where enrichment analyses report non-specific and non-robust findings, we are able to assess where we can be confident in their use. We find significant progress in recent bias correction methods for enrichment and provide our own software implementation. Our approach can be readily adapted to any pre-existing package.

## INTRODUCTION

As originally conceived, gene set analysis is a way to summarize rankings or groups of genes obtained from high-throughput experiments and as a tool for discovery (1–4). Broadly speaking, these methods look for statistical similarity between an experimentally derived gene set (or a ranked list of genes) and previously characterized gene sets (e.g., Gene Ontology GO (5), KEGG (6) or OMIM (7)). Running enrichment analysis on such data sets is now standard practice. Given the heavy reliance on these methods for hypothesis generation and experimental validation checks, it is important to improve our understanding of their benefits and limitations. As we will highlight, the central challenge in this analysis is how to manage and interpret results in light of gene set independence, or lack thereof.

One of the key insights into the challenge of gene set analysis is that some genes are simply generally more likely to be annotated to any sets. Such genes will appear in many sets. In the gene set analysis literature this property is often described in terms of overlap or annotation bias. In earlier work, we showed that the tendency for some genes to be frequently represented in GO is a critical confound in gene network analysis (8). A useful element of our approach in that work was to define redundancy within GO in terms of the ability of a single list of gene to predict the membership of each gene set derived from GO and its annotation. The degree to which a single list predicts all GO terms says how redundant GO is, which sets look to be most generic, and which genes contribute to those tendencies. Trivially, genes with many annotations would appear at the top of such a list because predicting them frequently will be correct across more GO groups. Thus, there is a strong overlap with “annotation” bias, but the two do have critical differences, as will be particularly evident when we assess approaches which correct for or minimize annotation bias. For simplicity, we refer to our calculation of the maximally predictive single list as estimating the gene’s “multifunctionality”, although the extent to which this form of multifunctionality represents a technical or true biological property will remain an open question.

In the gene set analysis context, because the redundancy and overlap in GO is often apparent when inspecting results, there have been a range of efforts to improve the situation (we use GO as our motivating example of an annotation scheme without loss of generality to alternatives). Many approaches attempt to reduce the redundancy in GO either by trimming it down up front (9–13), or adjusting the results of an analysis (14–16). An implicit understanding of the undesirability of overlaps of gene sets is also present when analyses are limited to a single branch of the GO hierarchy (e.g., only using Biological Process) or by using gene sets of a particular size range (e.g., less than 500 genes). Such approaches serve the dual purposes of simplifying interpretation of enrichment results and diminishing multiple test correction penalties, thereby improving p-values. However, attempts to reduce redundancy inevitably involve a loss of information, especially in schemes like GO where the extent of overlaps is extreme (8). Another approach to correct for redundancy is through improving the statistical machinery underlying gene set analysis, for example, by assuming that the underlying annotations are true enough to reconstruct the gene sets contributing to enrichment results by modelling their combined effects (17). More commonly, enrichment approaches make post-hoc adjustments, following a basic strategy of reducing the impact of multifunctional genes. Some approaches take the view that differential annotation for genes reflects a bias in the annotations that needs to be corrected, but that the correction needn’t depend on the experimental data on which gene set analysis is to be applied (18,19).

The commonality we point to in the various approaches is that it is hard to know if they improve upon what is already done. There is no strongly generalizable way to test the efficacy of these methods, as there are no gold standards. This is a problem likewise faced by any biologist in reading about and interpreting any results using any of these methods. But we take the stance that fixing the gene set analysis method or the gene set annotations is fraught with difficulties. Instead, our approach is akin to methods intended to test robustness (e.g., jackknifing) or overfitting (cross-validation), and is not a new form of enrichment analysis and thus can be applied to any gene set analysis method.

We rely on two central heuristics, uniqueness and robustness, which relate multifunctionality to the properties possessed by well-conditioned problems. Traditionally, well-conditioned problems are those that possess solutions unique and robust to minor data variation. For example, if enrichment output were identical to that produced by sets of genes that are present in many functions (i.e., multifunctional), then the results will not be at all uniquely characteristic. In such cases it will be hard to distinguish among the enriched functions which are returned; many functions will be returned and the distinction between the 100th function at p~1E-10 and the top function at p~1E-100 is not itself robust. Likewise, an enrichment result should not hinge on the presence or absence of any given single gene. Because we argue uniqueness and robustness are fundamental properties for the analysis to be meaningful, they will provide strong heuristic value to the interpretation of what would otherwise be a black box.

In this paper, we further develop and explore our model for enrichment and particularly the problem of multifunctionality, focusing both on detailed examples and a large corpus of studies. We derive ways to integratively assess multifunctionality as a confound that is applicable to multiple gene set analysis methods, including ones based on fixed thresholds (“hit lists”, e.g., (20)) and those which use complete rankings of all genes (e.g., GSEA (21)). We show that our approach improves the specificity of interpretation in enrichment analyses through an analysis across 17 commonly used enrichment methods. We propose that measurements of the effects of multifunctionality should be routinely incorporated in such analyses. To this end, we provide user-friendly implementations of the methods in a graphical user interface as part of the ErmineJ software package (22,23).

## MATERIAL AND METHODS

### Datasets

Except where noted, our analysis focuses on 20710 human genes, obtained using the UCSC GoldenPath (24) and NCBI databases (25). We downloaded the “C2” curated gene signatures from MolSigDB ([date: April 2013]) (26). We limited our analysis to 1800 lists of size 11-1000, which come from 659 different publications, and often form pairs (e.g., “up” and “down” regulated for the same experiment). Importantly, these lists are the type of “hit list” from genomics studies that typically forms the grist for performing GO enrichment analysis. In this paper we reserve the term “hit list” for such experimentally derived groups of genes to be analyzed, using “gene set” to refer to Gene Ontology GO groups.

### Gene Ontology and its derivatives

For the ErmineJ analyses, gene annotations were obtained via the NCBI Gene database (gene2go file [date: April 2013]), and the structure of GO was extracted from the XML files provided by the GO Consortium. As entailed by the semantics of GO, gene annotations were propagated to ancestors in the GO hierarchy based on is_a and part_of relations, excluding the roots (semantic closure expansion). We did not filter based on evidence codes and used all three domains of GO (similar results were obtained using just biological process or molecular function). Except where noted, we considered GO groups that had between 10 and 300 genes. There were 3172 GO terms meeting this criterion.

For subsequent enrichment analyses, we used the human gene association file (gene_association.goa_human, [date:12/12/2014]) downloaded from GO and the mouse version (gene_association.mgi [date:12/12/2014]). GO was constructed as above, using the OBO file (go.obo format-version: 1.2 [date:12/12/2014]).

We also constructed additional versions of GO using variations of the species (mouse and human), annotations (14 evidence codes), domains (cellular component, molecular function and biological process), and relations (only direct [omitting propagation], is_a, and part_of), each of which can be used to provide gene sets for gene set analysis. Using every pairwise combination of a single choice of each property, including “all” and “none”, over all genes or only those jointly present, yields 512 GO and annotation combinations. For each of these derivative GOs, we calculated the gene multifunctionality scores (see below) and assessed the fraction of GO terms enriched on this list at an FDR<0.05 and FDR<1E-10.

Further to parsing the role of properties within the existing GO and its annotations, we generated four novel versions of GO (alto-GOs), encompassing an alternate conceptualization of how an ontology, annotations to it, and methods exploiting the two, interact. We labelled these Shadow-GO, Ortho-GO, Weigh-GO and Local-GO (see Supplement for more details), collectively termed “alt-GOs”. For Shadow-GO, each GO term brings into existence its complement to which genes would be annotated as “not” being members. This additional “Shadow” of the original GO would follow valid rules of inference defined by GO through *modus tollens*, and all genes have an identically equivalent number of annotations. In this alt-GO, the annotations are changed, but the ontology is the same. Ortho-GO alters GO by performing dimension reduction on the original matrix of propagated GO annotations, yielding new genes sets which are closer to independent but retain the original tendencies of pairs of genes to be co-annotated. Weigh-GO discards binary membership, such that each gene is weighted based on set annotation specificity. Local-GO is more targeted version of GO, where we select a function of interest, and pick non-overlapping GO terms to also test. In this case, the annotation sets are held constant, but the ontology is tweaked to only include a subset of groups. To generate and assess this, we pick a random function within GO to be of interest and then iteratively pick new functions based on the minimum Jaccard overlap with the remainder, stopping at either 200 or 1000 local functions (200-local-GO and 1000-local-GO, respectively).

### Multifunctionality of genes

We describe multifunctionality in terms of the number of annotated GO terms for the gene our case studies but use the analytic calculation from (8) throughout:

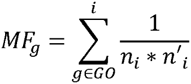

where *i* are the GO groups that gene *g* is a member of, *n*_*i*_ the number of genes in that group (in the universe of all genes), and *n*’_*i*_ those not in the group (the complement). The final score is then ranked and standardized. The equation takes this form as it is the ranking that maximizes the area under the receiver operating characteristic curve (AUROC) for each gene set under consideration, averaged across all gene sets (8). In the real GO it is highly correlated (r>0.95) with the number of gene terms a GO group has (being a version of this weighted by specificity), and our results are generally robust to either choice, except where methods attempt to specifically correct away annotation bias, as in the hypothetical ontologies.

### Enrichment analysis

We considered two basic types of algorithms. First is a “basic” enrichment analysis based on the hypergeometric distribution, and which requires defining a “hit list”. The second is based on ranks without setting a threshold. For this purpose we used a method based on the AUROC (27), the same as the method mentioned above to measure multifunctionality of a GO group but using the experimentally-derived ranking. ErmineJ implements several additional methods, including the resampling methods described by (2) and a GSEA-inspired method that uses precision-recall analyses rather than modified Kolmogorov-Smirnov statistics, in which the mean average precision (similar to the area under the precision-recall curve) is calibrated by random sampling to obtain a null distribution. The false discovery rate (FDR) was controlled using the method of Benjamini and Hochberg (28).

### Multifunctionality analysis of enrichment

For the hypergeometric method, we take the approach of testing the effect of iteratively removing genes from the “hit list” in order of multifunctionality. The challenge is identifying an appropriate stopping point. Our algorithm is motivated by finding a point at which the enrichment results are maximally sensitive to the removal of the most multifunctional gene. Intuitively, if some gene sets are only enriched due to overlaps, as we remove overlapping genes, those gene sets will eventually fall away. This transition point will be reflected by a rapid alteration in the most significantly related gene sets, similar to the phenomenon shown in **Figure 3C**. We found this is effective in finding the optimum stopping point in model data, and is also of value as a test for “robustness” in enrichment. A formal description of the gene removal algorithm and a schematic is given in the supplement (**Supplementary Figure 1**). If the hit list is not significantly multifunctionality-biased (based on the Mann-Whitney U test as described for GO groups above, p<0.05), or if no gene sets are significantly enriched at a pre-set FDR q (we used q=0.05), no correction is performed. If the algorithm iterates such that more than ½ of the genes in the ‘”hit list” are removed, the algorithm terminates.

For methods that use a full ranking of genes, we developed an approach using regression. For the ROC-based method (27), the appropriate regression was unweighted linear regression of the genes scores against the gene multifunctionality scores; the original gene scores are replaced by the Studentized residuals of this regression. Thus genes which are highly ranked, but also highly multifunctional, will tend to be “bumped down” in the ranking. We note that some methods, such as GSEA, use the full ranking but behave more like precision-recall curves than ROCs, in that they put much more emphasis on highly ranked genes. In this situation unweighted regression is inappropriate. While not investigated as part of our analysis reported here, a regression-based correction for the precision-recall method is implemented in ErmineJ 3.0. The regression is weighted by 1/√N where N is the rank. This can be motivated by observing that under the null distribution of random rankings, the standard error of the precision is higher at low recall, and this is expected to vary as 1/√N (making the simplifying assumption of independence of the genes). This variability is what determines the expected contribution of a hit to the aggregate variability in the area under the precision-recall curve.

### Disease association analysis

Disease-gene relationships, organized by disease ontology (DO) terms, were obtained from Phenocarta (29) [date: April 2013]. Enrichment of MolSigDB hit lists for these disease gene groups were performed using ErmineJ 3.0. MolSigDB hit lists were associated with DO terms using the National Center for Biomedical Ontology (NCBO) Annotator (30), applied to the title, abstract and Medical Subject Headings (MeSH) associated with the linked PubMed record for the MolSigDB list.

### ErmineJ implementation and analysis of case studies

ErmineJ implements multifunctionality analysis as well as the unweighted and weighted regression correction algorithms. ErmineJ implements gene multifunctionality as defined by (8), as well as reporting the simpler “number of annotations” measure. For the case studies reported here, ErmineJ analyses were limited to the biological process GO aspect, for terms containing 20-200 genes. Gene lists for case studies were extracted from data presented in the original reports or supplementary tables. The gene lists we discuss are based on the identifiers we could match to official gene symbols in our database, so may not exactly match the lists reported by the authors. The data files used for the case studies are available in the online supplement (http://ErmineJ.chibi.ubc.ca/multifuncsupplement/).

### Analysis of alternate enrichment methods

We selected 17 common methods that perform varying forms of gene set enrichment and correction procedures (accessed between Dec 2014 and April 2015). For the most part, these methods rely on a statistical test to determine which gene sets are significant and some method of enrichment correction. Here we focus on methods specifically designed for GO. We ran each method with the same default parameters, and when we could, used the same background input. The GO annotations file also varied as some methods had set their annotation file, and others allowed the user to specify it. For consistency, we attempted to use the same GO version when possible. Because we could not directly control for the number of GO terms used, we attempted to control for this by comparing the fraction of GO terms returned, instead of totals. However, we did not wish to penalize methods, so we continued to compare all results, even if some GO terms were missing between methods. The total set of GO terms with gene annotations was then almost 16K, and for each case study methods reported between 2000 and 6000 terms. We did not limit the results to a particular GO category and excluded IEA annotations as is commonly done for purely algorithmic assessments, since IEA annotations are themselves algorithmically determined.

To calculate the functions different methods are likely to return for most hit lists, we performed GO enrichment analysis, using a list of the top 100 multifunctional genes derived from human GO annotations as of Dec 2014, for the 17 different methods, and variations of a few of these methods, including ErmineJ and a basic gene set enrichment implementation (hypergeometric test). We calculated the number of GO terms returned as significant for this list of 100 genes and how these terms and their p-values correlated between methods. We chose a p-value threshold of 0.05 to compare the number of results returned by each method. Some methods return all tested values, while others only the significant terms they found enriched. Most methods perform their own multiple hypothesis test corrections, and when able, we specified for Benjamini-Hochberg. All these analyses were similarly repeated for mouse GO annotations.

For the uniqueness assessment, we took each case study and compared the enrichment results to the multifunctionality results, first by calculating the average multifunctionality of the GO terms returned as enriched, and also comparing the overlap of results from the previous multifunctionality enrichment, species specific. We then performed a robustness analysis using those same case studies. For this, we removed 5% and then 10% of the most multifunctional genes from the list, and re-ran the individual enrichment methods. We then calculated the overlap between the enrichment results returned for each method, as a measure of stability. We also then once again compared how multifunctional the results were once we removed the most multifunctional genes.

Additional information including data files for many of the analyses and scripts are available online at http://ErmineJ.chibi.ubc.ca/multifuncsupplement/.

## RESULTS

In this paper, we evaluate the effect of multifunctional genes on enrichment results. We start by outlining our motivation and illustrating the impact of multifunctional genes in our model of uniqueness and robustness. We move on to providing specific examples in four case studies. We next perform an assessment of multifunctional genes in globally used gene lists using our standard algorithm. We then demonstrate the impact of multifunctionality bias across multiple algorithms and extend our multifunctionality analysis to alternate versions of GO. We conclude with a demonstration of the ErmineJ software where we have implemented the multifunctionality bias assessment.

### The multifunctionality problem in gene set analysis

To illustrate our work’s motivation, we outline a conceptual model for gene set analysis that characterizes a specific experimental outcome (**Figure 1**). In the model, the system being studied is presumed to involve several gene-based “functions”, which together contribute to the observed cellular or physiological state (e.g., disease or phenotype.). With some probability, genes in those functions will be detected in the study (depending on the type of assay, the strength of the signal, etc.). The more multifunctional a gene is, the higher the prior probability of it possessing any given function. If existing gene annotations (e.g., GO (5) with its annotations (31)) capture the relationships of the detected genes to those functions, we would expect an enrichment analysis to rank those functions highly. We note this model uses the “competitive” null hypothesis in the framework of (32), in which genes with an annotation are contrasted with those that do not.

**Figure 1.**
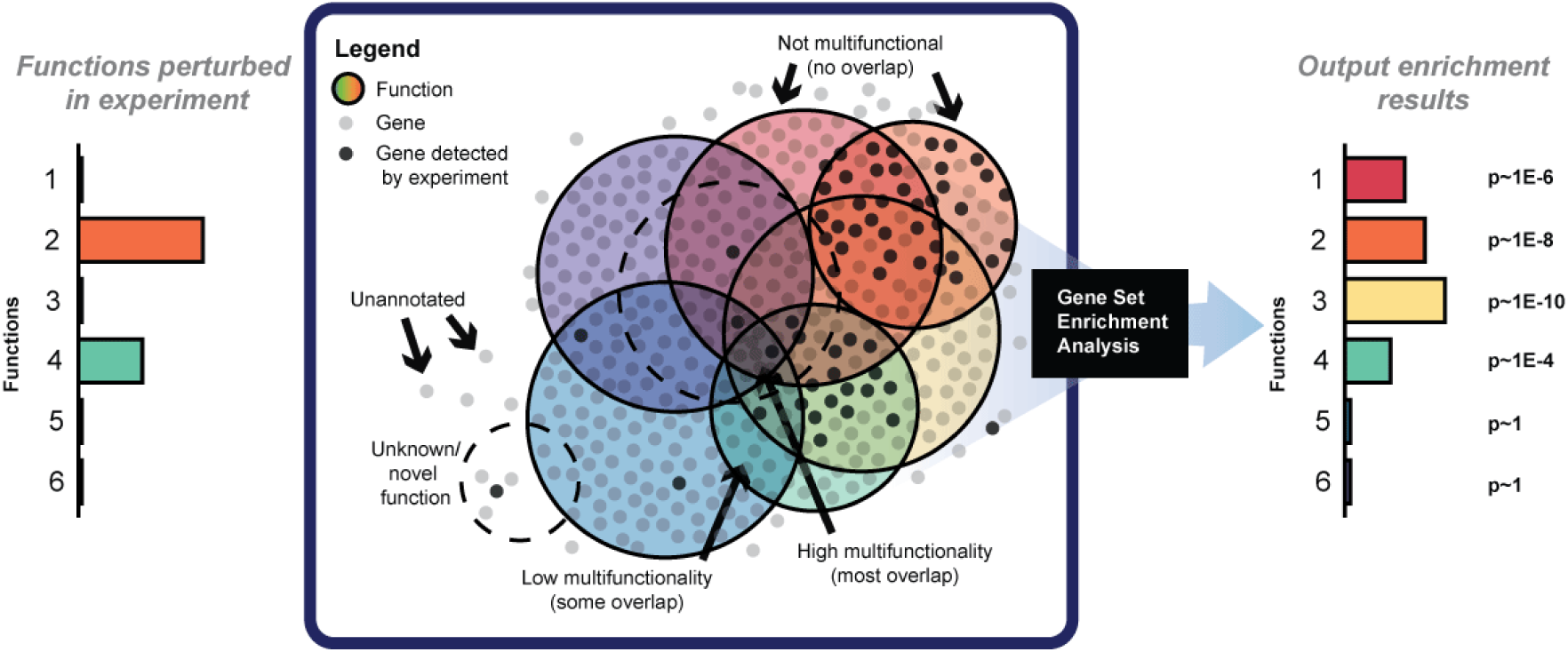
A conceptual model for gene group analysis. Consider a hypothetical process (e.g., a disease) which involves two gene functions (bars at left). Assume a gene is “detected” in the experiment (with some non-zero probability) if they are involved in one of the functions that underlie the process of interest. In this case, the genes with the highest probability of “showing up” are the ones in functions 2 and 4. From a gene set enrichment analysis of this hit list, it is hoped that enrichment will be found for both functions 2 and 4, but not the others. Yet, genes in these functions are highly multifunctional and share other functions, which show up erroneously as enriched in the analysis (bars on the right).

Ideally the enrichment results will reconstruct the underlying functions. Because genes commonly have more than one function (i.e., are annotated to more than one GO group, pathway etc.), it is possible that the group which is most enriched is not actually one of the functions that is truly involved in the process or phenotype. This can occur if a group preferentially contains these “multifunctional” genes which overlap with the genes annotated to the true functions. Since many statistical analyses assume all genes are equally likely to be perturbed in the experiment, enrichment analysis here could be highly misleading. For example, if we truly believed two functions were involved in some experiment, then we would predict the experimental outcome (gene sets enriched) to most closely resemble not the “input functions” individually, but whichever third function is best characterized by their overlap. The situation, which we have summarized in **Figure 1**, is extremely simplified, as real data is far more complex, with hundreds or thousands of groups and many opportunities for such overlaps to occur. And, of course, when the multifunctional genes at the intersect are enriched, many functions will appear as ‘significant’ without being meaningful.

### Tests for multifunctional effects in gene set enrichment

To interpret gene set enrichment results in the light of multifunctional genes, we suggest a series of tests that can be applied to the output of any analysis. As previously mentioned, well-conditioned problems are those that possess unique solutions and are robust to data variation. We demonstrate the effects of multifunctionality on these uniqueness and robustness heuristics in **Figure 2**. As a demonstration of the first test, imagine the input to a gene set analysis was the genes ranked by the number of GO functions they possess (we use this list repeatedly in this paper, referring to it as the “gene multifunctionality ranking”, with the most heavily-annotated gene at the top). Applying a simple enrichment test to this ranking (Mann-Whitney) we find 92% of GO groups (with at least 5 genes) are significantly enriched (FDR<0.05), and 36% at FDR<1E-10. When we use the multifunctionality ranking as described in previous work that takes into account GO set sizes along with number of GO terms (8), we find even greater enrichments, with 98% at 0.05 and 36% at 1E-10. The degree to which the actual input to a gene set analysis has any resemblance to the multifunctionality ranking will result in a concomitant similarity to the biologically non-specific (non-unique) results of enrichment analysis of the multifunctionality ranking. Thus, using the multifunctionality ranking as a comparator to the actual ranking provided as input will demonstrate the uniqueness of the output. To further elaborate, in the case of experiments that return multifunctional genes in their hit lists, it is possible to get similar if not identical results from an enrichment analysis (**Figure 2A**). Genes in the hit lists may not be identical between the two experiments, but they share common functions, and the resulting enrichment will therefore be non-specific and non-unique.

**Figure 2.**
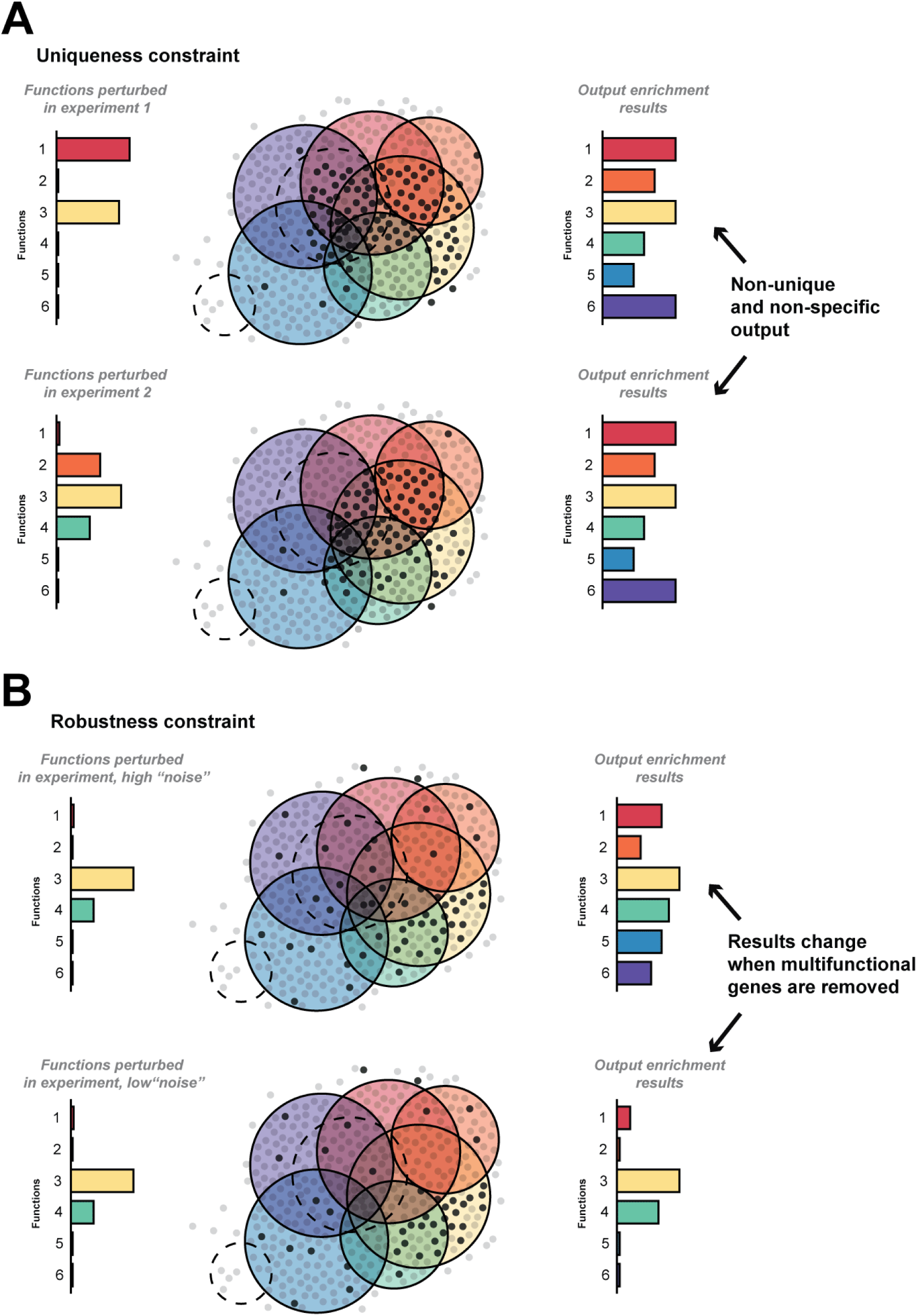
Uniqueness and robustness as constraints for testing validity of enrichment outputs. (A) For a given experiment 1 with unknown (or known) functions 1 and 3 (bar on right in top panel), an experiment detects a set of genes (marked in dark grey in figure). Gene set enrichment outputs results (bar on left in top panel). Because the genes detected are multifunctional, many functions are returned enriched. For a different experiment with a different phenotype or disease, the functions (right bar in bottom part of panel), are different (or overlapping), and the genes detected may be similar. Enrichment yields similar if not identical results to the previous experiment due to the multifunctional genes. Comparing this output to that of the multifunctionality assessment shows that the results enriched are non-unique. (B) When testing robustness, removing genes such as the most multifunctional has little impact on the non-unique results, but removing genes from functions in experiments with less “noise” (i.e., fewer overlaps that are multifunctional), removes the non-specific signals.

Multifunctional effects are critical to the assessment of robustness as well. For an experiment that perturbs genes with many functions, the results will be robust to variation – removing multifunctional genes will likely keep these non-unique results. In the case of an experiment with fewer overlapping perturbed functions, removing the multifunctional genes will remove results that are due to the multifunctional effects, leaving behind a more robust set of functions (**Figure 2B**). Removing the most multifunctional genes will impact the overall output, and thus the stability of the enrichment can be used as gauge. If removing a single gene can disrupt the results, the point at which doing so has little effect, is the point where we can be more confident in a biological interpretation of the output. This idea is the basis of the multifunctionality correction we suggest in our algorithm (detailed in the supplement). We go into more detail in the following sections on these points, applying these tests to specific case studies and global gene lists.

### Characterizing multifunctionality effects in case studies

Before presenting the details of the algorithm we developed for multifunctionality assessment, we first describe the results the assessment of multifunctional genes provides for four motivating case studies. While these case studies were selected because they show interesting effects of multifunctionality, they are far from being unusual, as we describe in later sections. Each case study is based on the list of genes identified by the investigators as being of interest. We refer to these as “hit lists” to differentiate them from the gene sets which are tested for enrichment. We used ErmineJ as the algorithm for gene set enrichment. We refer to multifunctionality-corrected results as those obtained by removing the most multifunctional genes in the hit list, which serves as the basis of the robustness tests in our algorithm. More details are included in the Methods section and data and full results files for each case study are presented in the online supplement.

#### Genomic copy number variants in autism

Based in part on GO enrichment analysis, Gilman et al. (33) hypothesized that synaptic development and function is at the heart of the autistic phenotype. Repeating their analysis using ErmineJ on their hit list of 70 genes, we arrive at a list of 30 GO gene sets (FDR 0.05) similar but not identical to those reported. Assessing the multifunctionality of these 70 genes, we see that the list is quite strongly biased towards multifunctional genes (MF score=0.86, p<1E-8). Removing the 11 most multifunctional genes (*FLNA, NRXN1, DLG1, MAPK3, CRHR1, DLG4, DKK1, AXIN1, WNT3, NLGN3, STUB1*) has an important influence on the results and their subsequent interpretation. Firstly, the heavy down-weighting of the “learning or memory” GO gene set provides a good illustration of the benefit of considering multifunctionality (**Supplementary Table 1, Supplementary Figure 7**). Of the six genes in the cluster that have this annotation, four are among those 11 reported as highly multifunctional (*NRXN1, NLGN3, DLG4, CRHR1*); they have between 186 and 289 GO annotations each. By down-weighting such genes, weaker signals were allowed to be more prominent. For example, the term “neuron migration” (supported by four genes among the 70), was originally ranked 28th in our analysis but is unaffected by multifunctionality correction and thus rises in the ranks. From a biological perspective, neuron migration might be even more relevant to ASD than learning and memory. However, we stress that the enrichment of “learning and memory” in the first place is not a statistical false positive; we prefer to think of it as non-robust and, most importantly, non-specific. The importance of *NRXN1* and *NLGN3* to ASD was already bolstered by a simple analysis of the genes contained in the CNVs studied by Gilman et al. (34), and their heavy annotation helps ensure that they drive the enrichment results.

#### Genome-wide association studies of schizophrenia

Schmidt-Kastner et al. (35) identified 77 schizophrenia (SZ) candidate genes from a review of the genetic association literature, and used enrichment analysis and manual annotation to gain support for their hypothesis on vascular stress responses. Our re-analysis of this list shows it is also strongly multifunctionality-biased (p<1E-14), with GO enrichment results related to stress and inflammation. However, a multifunctionality-corrected analysis results in no gene sets meeting the significance criterion, yet those at top of the list now involving synaptic transmission. The two points illustrated by this case are the dependency of significance on the multifunctionality of genes as none pass multiple test correction, and it is often possible to construct a variety of narratives when faced with a multifunctional gene list. In this case, the same list of genes could be treated as having something to do with synaptic transmission, while from another point of view it has something to do with stress responses.

#### Gene expression changes in response to hypoxia

Manalo et al. (36) identified genes which were changed in RNA expression in response to hypoxia, and intersected with genes which were induced by a constitutively active form of the HIF-1 transcription factor. As for the other case studies, their list of 202 genes was biased towards multifunctional genes (p<1E-10). Unsurprisingly, the enrichment results were also highly sensitive to removal of multifunctional genes, as after multifunctionality correction, only four groups would be significant at an FDR of 0.05 (70 originally). These include “peptidyl-proline modification”, “cellular response to hypoxia”, “collagen fibril organization” and “cellular response to oxygen levels“, which strongly align with the themes the authors chose to focus upon. We argue that this “cleaning up” of the results increased their relevance to the study while not precluding the investigation of less specific terms.

#### Protein interactions of Oct4

Pardo et al. (37) studied mouse genes whose products were found to physically interact with the Pou5f1 transcription factor (more commonly known as *Oct4*), a crucial protein in the regulation of cellular differentiation and thus in embryonic development. In ErmineJ, the list of *Oct4* interactors yields enrichment of many GO gene sets related to DNA and chromatin function, spanning recombination, replication and histone acetylation, but also a variety of other processes such as “modification of symbiont morphology or physiology” and “ATP metabolic process” as well as terms relating to embryonic development. A multifunctionality-corrected analysis yields a shorter and more focused set of gene sets which emphasize chromatin remodeling and histone acetylation. Importantly, removing *Oct4* from the list of interactors dramatically reduces the number of GO gene sets considered significant after multifunctionality correction, leaving just one, “chromatin remodeling”. This illustrates the impact even a single highly multifunctional gene can have on an enrichment analysis and reiterates how protein interactions are highly biased towards multifunctional genes (8,38).

### Enrichment results are sensitive to gene multifunctionality

Having characterized the effects of multifunctional genes in the individual case studies, we wished to obtain more insight into whether multifunctionality might affect enrichment analyses globally. We next examined a large set of experimentally-derived gene lists, primarily from transcriptome profiling studies (MolSigDb, (26)). We evaluated the multifunctionality of genes within these hit lists, and their impacts through simulation studies and the robustness and uniqueness tests.

#### Multifunctional genes are overrepresented in genomics results

**Figure 3A** shows that genes which show up on multiple MolSigDb lists tend to be multifunctional (Spearman’s rank correlation r_s_= 0.48). This suggests that multifunctionality defined by GO is reflected in the responses of genes to experimental manipulations. In other words, genes which are highly annotated tend to turn up more often in genomics studies. Whether this is a cause or an effect of the annotations, is not entirely clear (8). For example the trend in **Figure 3A** is likely confounded by biases in the choice of which genes are analyzed in each study; most studies in MolSigDb used microarrays that contain probes corresponding to many but not necessarily all known protein coding genes, and genes which are “popular” are more likely to be tested. Regardless, this analysis supports the view that multifunctionality measured by GO is relevant to the analysis of genomics studies.

**Figure 3.**
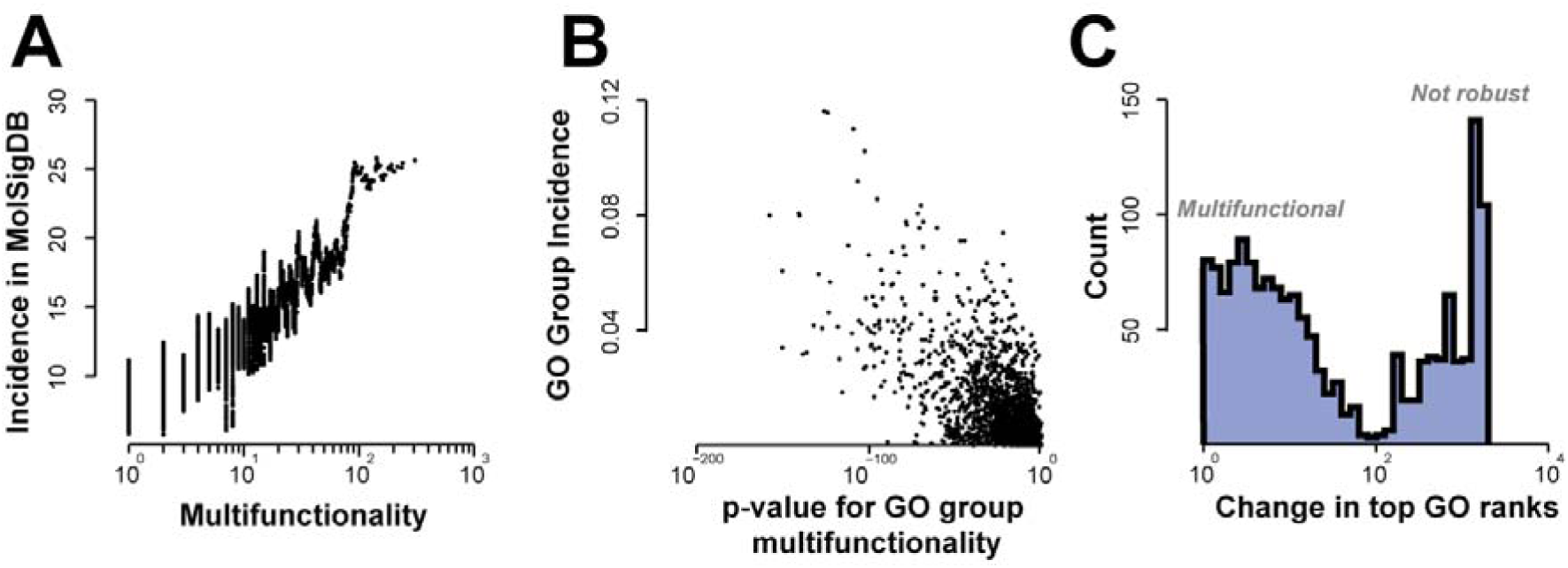
Multifunctionality strongly impacts GO enrichment results on published “hit lists”. (A) Multifunctional genes appear more often in MolSigDb lists. Multifunctionality is the number of GO terms assigned to a gene. Data are smoothed with a sliding window of 100 genes. (B) Multifunctional GO terms are more frequently enriched in MolSigDb hit lists. P-values for AUROCs for GO functions using gene multifunctionality ranking versus GO group incidence results of simple over-representation enrichment analyses of MolSigDb groups (threshold FDR<0.05). The Pearson correlation is r=-0.67 using the log p-values as shown, Spearman’s rank correlation r_s_=-0.59. (C) Enrichment results are sensitive to the removal of single genes depending on multifunctionality of the GO term. Change in top 10 GO ranks for each hit list after removing one gene (the strongest contributing one). Multifunctionality-enriched hit lists tend to gather at the left; (at a FDR<0.05, mean shift is 8) whereas the sets that are not enriched for multifunctional genes change by an average of 902 (right-hand peak).

Given the trend in **Figure 3A**, it is not surprising that we observe a similar phenomenon for GO term enrichment, in which gene sets found to be enriched in multiple MolSigDb lists tend to be enriched for multifunctional genes (**Figure 3B**; Spearman’s rank correlation r_s_= -0.67; distributions of multifunctionality scores for GO and other gene set schemes are shown in **Supplementary Figure 2**). That is, gene sets defined by GO which contain genes which are multifunctional (have many GO terms) are more likely to be enriched in the MolSigDb lists. Such gene sets, by virtue of their highly annotated members, will tend to be less biologically specific. A related evaluation using MeSH terms is shown in the supplement (**Supplementary Figure 3**).

We next performed a type of sensitivity analysis, where we tested the impact of each gene in a hit-list on the results of the enrichment analysis. We find that for many MolSigDb lists, the enrichment results are highly dependent on the presence of one gene (right-hand peak in **Figure 3C**). The gene which causes the largest shift in the results was preferentially the most multifunctional gene (r=0.35). In another group of lists, the results were insensitive to the removal of any one gene (left-hand peak in **Figure 3C**). These hit lists are found to be the ones which had multiple multifunctional genes, rendering the removal of any one gene ineffective.

Taken together, these results suggest that the outcome of a gene set enrichment analysis can be highly dependent on the presence or absence of multifunctional genes in the “hit list”. Further, the gene sets that are found by enrichment analysis tend to be multifunctional, thus having less specific interpretations. This strongly suggests that the presence or absence of multifunctional genes will be informative in determining whether results from an enrichment analysis can be trusted.

#### Exploring multifunctionality through simulation studies

Our first model, designed to test the effect of multifunctionality in a relatively simple case, examines a hypothetical experiment that yields a “hit list” of 100 genes, to which we wish to apply a hypergeometric test to evaluate enrichment. Ten of the genes in the list come from one target GO gene set (that is, annotated with a particular GO term). The other 90 are assumed to be irrelevant noise. Ideally the target GO gene set should rank very highly, if not first, in the enrichment analysis. Indeed this will usually happen if the 90 other genes are chosen completely randomly. However, if we add an additional constraint that the 90 genes must have a minimum degree of multifunctionality (but still selected at random), the situation changes dramatically (**Figure 4A**). If the “background” genes are too multifunctional, the target GO gene set is no longer successfully retrieved. This is a direct demonstration of the **Figure 1** scenario.

**Figure 4.**
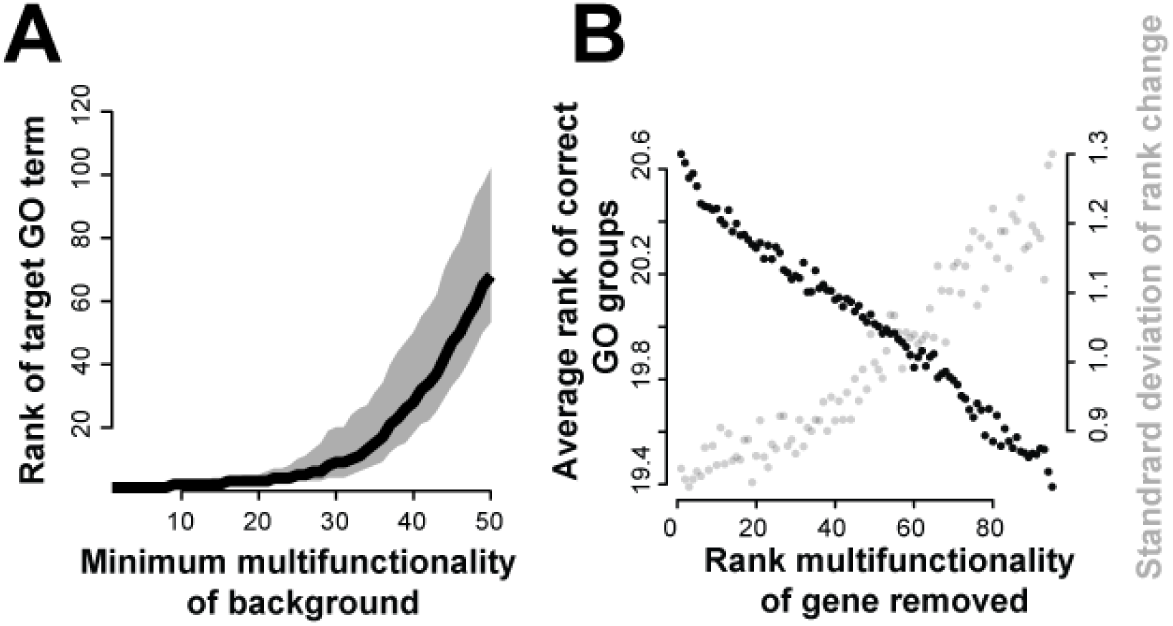
Simulations showing that the presence of multifunctional genes degrades recovery of “true” functions. (A) Analysis of a simple simulation, in which hit lists of size 100 are generated using 10 genes from a randomly selected GO group (which becomes the target) plus background noise of 90 randomly chosen genes, with the background constrained to have a minimum multifunctionality level. Increasing the minimum multifunctionality of the background genes (x-axis) decreases recovery of the target GO term. Black line indicates average of 1000 simulations; grey area covers 50% of simulations (middle quartiles). (B) Analysis of a more complex model, mixing 10 genes from 10 GO groups to make an artificial hit list, and testing the effect of removing each gene (one at a time, not cumulatively). We plotted the average rank of the target functions against the multifunctionality rank of a single gene removed. Because only a single gene is removed, the effect is modest, but the more multifunctional the gene, the more removing it improves the score but also the more the scores vary. Note the y-axis only includes the range from 19.3-20.7. The plot is the average of 1000 simulations, each of which involved removing each of the 100 genes in turn.

We next explored a more complex model, where ten GO gene sets were randomly selected as targets, and ten genes randomly selected from each (because a gene could appear in more than one GO gene set, this sometimes resulted in slightly fewer than 100 genes in total). The ideal enrichment analysis result would be that the top 10 GO terms would, on average, be the ones which were used to construct the hit list, yielding a mean rank of 5.5. Simulating this situation 1000 times, the mean rank of the ten target GO gene sets was 20.7. We then removed a single gene from the hit list and repeated the analysis, doing this for each gene in turn, for each simulated set (1000 × ~100 simulations). As shown in **Figure 4B**, the improvement in the result is proportional to the multifunctionality of the removed gene. That is, removing the most multifunctional gene has the tendency to “clean up” the enrichment results so the truly underlying functions are closer to the top of the ranking (the theoretical optimum of 5.5 is not attainable in this simulation due to overlaps among GO groups). An interesting aspect of this result is that the gene being removed is a “true positive”, in the sense that it belongs to at least one of the 10 target gene sets. Thus even though the enrichment signal for groups it belongs to is necessarily weakened by its removal, the cost incurred by including it is even worse, owing to its multifunctionality. However, this improvement in specificity comes with a cost in the form of far higher variation in correct rankings when the multifunctional gene is removed (**Figure 4B**, grey). This suggests that removal of multifunctional genes will both improve specificity and reveal underlying fragility in results by determining the potentially erroneous feature on which they most critically depend. Further simulations and results are shown in **Supplementary Figure 4 and 5** and the comparable results for continuous corrections are shown in **Supplementary Figure 6**.

#### Field-wide evaluations for the impact of multifunctionality

While our simulations were useful in framing the problem with multifunctionality, it is obviously crucial that the approach (i.e., evaluating and/or removing multifunctional genes) has desirable effects on real data overall. This is difficult to evaluate because in real data there is no established gold standard for enrichment results. This has been a consistent challenge for all such evaluations in the literature. As we will not know what the functions a hit list should be enriched for, we perform our next analysis once again on the standard MolSigDB hit lists, and identify the impact of multifunctional genes on what would be reported.

We wish to detect the point at which reported results are both robust and unique to removing multifunctional genes; in other words, the point at which overlaps have decreased and the influence of individual genes is small. Since we found that multifunctional genes most contribute to large swings in enrichment, this argues in favor of removing multifunctional genes in descending order until reported results are no longer sensitive to their removal. If we remove multifunctional genes in descending order, we also slowly remove functions from the reported enriched results. In our case, this resulted in 8% of genes being removed on average, with 52% fewer GO gene sets being significant (at an FDR 0.05). Note that this does not yield additional enriched functions, but merely argues that a subset of apparently significant results were not robust (and primarily due to high prevalence genes). More importantly, there is a decrease in the occurrence of enrichment of highly multifunctional GO gene sets. The correlation between occurrence and multifunctionality goes from r=-0.67 (as in **Figure 3B**; Pearson correlation) to r=-0.51. Further, previously rare gene sets became more common, leading to a more even spread of occurrence of GO gene sets across the MolSigDB results (the standard deviation of times a term occurs in the results goes from 24 to 13.3 after this ‘correction’). The agreement of these results with the expectations under the model (**Figure 1**) supports the hypothesis that over-occurrence of multifunctional groups is an artifact, not a meaningful biological phenomenon. More importantly, it suggests that multifunctional genes are a good place to look to for irreproducible results. If removing a single gene – and particularly the gene which was likeliest to arise by chance anyway - removes most of your enrichment, the enrichment results were probably not reliable in the first place.

### Characterizing algorithms using the robustness and uniqueness heuristic

Having assessed the impact of multifunctionality in a baseline algorithm, we now look to see if the same ideas apply to other methods. Our focus here is on demonstrating that simple assessments of multifunctionality reveal more clearly what pre-existing methods are doing. We once again consider the four case studies previously described, but now consider a corpus of 17 pre-existing enrichment analysis methods (including our own ErmineJ). We use all the methods as black boxes, using their default settings but standardizing so that the inputs and outputs are comparable.

Recall that using the gene multifunctionality ranking as input to a simple Mann-Whitney enrichment test yields over 90% of gene groups enriched. The extent to which the results of an actual analysis resemble this result is a measure of interest in the uniqueness analysis (**Figure 5A**), which is naturally customized to each algorithm since the exact functional outputs in response to this generic input will vary. In response to an input of the top 100 multifunctional genes as an input (see methods), the methods returned an average of 22.38% (0.12%-77.44%, SD 22.25%) of all GO functions as significant, indicating the functionally promiscuous effect even this small set of genes can have in some methods. The wide range indicates that methods vary strongly but a substantial fraction of that variance may be enforced by direct filtering (GO groups which can’t be returned). This highlights the importance of having a way of benchmarking the outputs of real enrichment gene lists since variation in output for a given method will reflect the same filtering. The precise p-values for GO terms differ from method to method, with correlations of p-values across significantly enriched terms averaging r_s_=0.30 (Spearman’s rank correlation, SD 0.21, **Figure 5B**).

**Figure 5.**
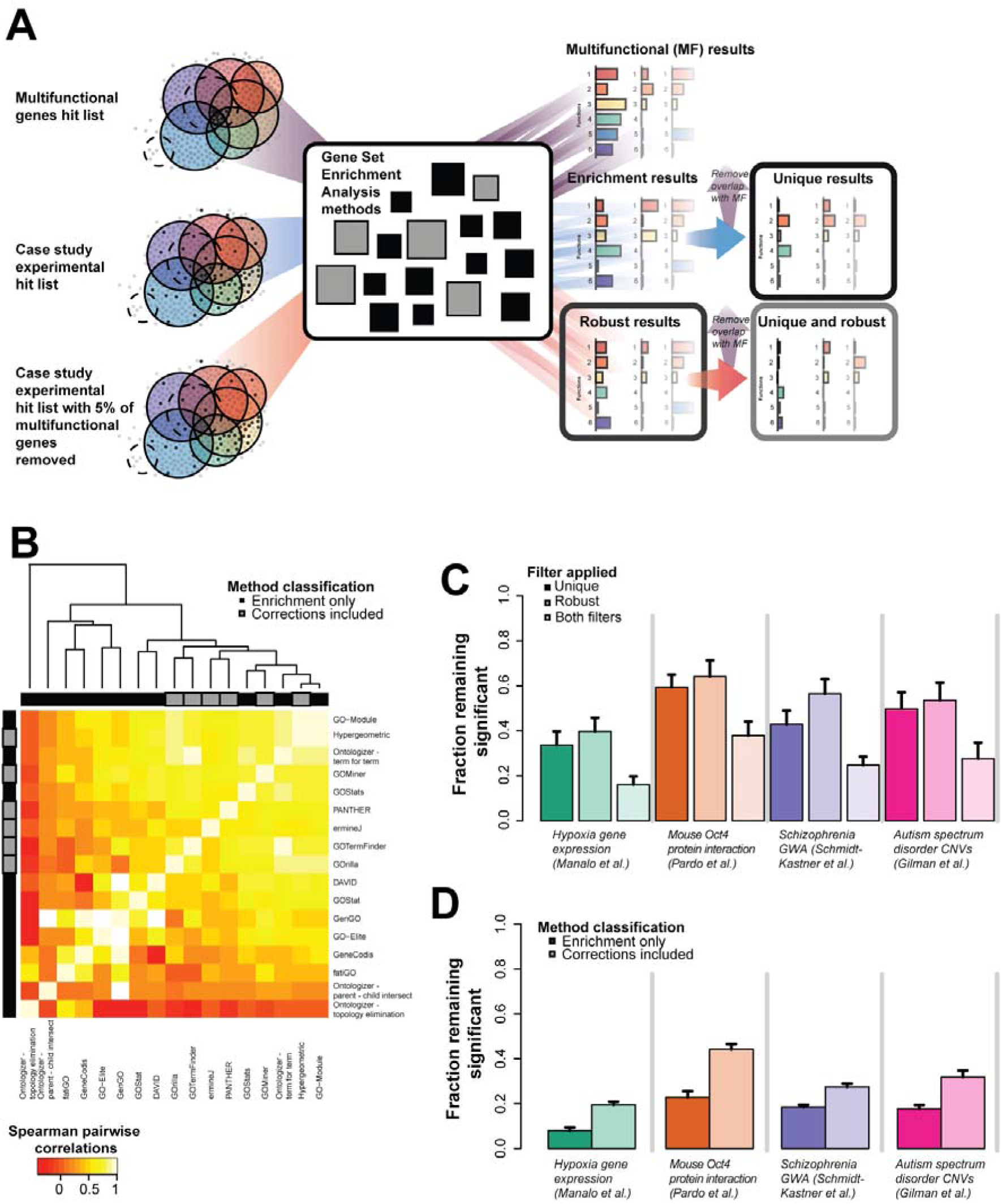
Effects of multifunctionality on algorithm behavior. (A) Schematic of method to assess uniqueness and robustness of the 17 gene set enrichment methods. We input the top 100 multifunctional genes, the case study genes, and then the case study genes filtered at 5% for the most multifunctional genes. The 5% reduced output results are those robust to multifunctionality. The results filtered by the multifunctional results are those used in uniqueness test. (B) The top 100 multifunctional genes were given as a hit list for the individual algorithms, and the resulting GO enrichment results for each were compared. The methods that do not claim to correct cluster together. The corrections that prune results post-enrichment cluster with the non-correcting methods. (C) Four case studies were assessed in each of 17 commonly used enrichment methods and their results assessed for the role of multifunctional genes in generating their systemic results. Only a modest fraction (average 46.4%, average SE 6%) of the reportedly enriched functions are not the same ones that each algorithm outputs when the 100 most multifunctional genes are used as an input (leftmost panels). Removing the 5% most multifunctional genes from each hit list (as few as 1 gene) dramatically alters most reported enrichment, leaving only ~53.5% (average SE 7%) of them in tact (middle panel). This combination of effects has an impact on all but a small fraction (average 26.6%, average SE 5%) of the algorithms across all four studies (right most panels). Note that colors associated with the study are indicated in the legend in panel B. (D) Algorithm behavior is examined for the effects of corrections. We partitioned the algorithms into two classes, those which perform more standard statistical tests (darker colors) and those which attempt to correct for problems with enrichment in some way (lighter colors). We then repeated the analysis from part A. Algorithms attempting to correct their output yield a significantly higher fraction of terms which are both specific (not multifunctional) and robust (to removal of 5% of genes from the hit list).

We now apply our criteria of uniqueness and robustness to the output of the 17 algorithms for our four case study data sets. First, to assess uniqueness, we compared the output of each algorithm when given the experimental input hit lists to that of the algorithm when the top 100 multifunctional genes was the input (**Figure 5A**). Recall that the multifunctional hit lists just input genes with as many functions as possible, so output enriched functions may be significant but appear only to the extent they overlap with other functions, and are therefore non-specific (as illustrated earlier in **Figure 2A**). Because we know the four case study data sets have a substantial multifunctionality bias, as expected the overlap in the enrichment results with the top 100 multifunctional genes and the experimental hit lists is very high. Filtering out this overlap results in an average retention of only 46.4% of the results, SD 10.8% (darker shaded bars in **Figure 5C**).

We next assessed robustness by removing the 5% of most multifunctional genes from the experimental hit lists (as demonstrated in **Figure 2B** and **Figure 5A**). While this percentage is arbitrary, it seems an extremely conservative test to us (if removal of only 5% of the hit list – as little as one gene – can alter reported enrichments, it would seem unreasonable to consider the results meaningfully robust). As for the comparison with the top 100 multifunctional genes, we compare the results after this removal to the original results with the hit list, finding an average of ~53.5% (SD 10.1%) of reported enrichments are retained (lighter shaded bars in **Figure 5C**). This confirms that many enrichment results in these case studies are not robust.

We next combined the uniqueness and robustness filters, yielding an average 26.6% (SD 9%) of GO terms robust to both (lightest shaded bars in **Figure 5C**). It may seem surprising that the filters are not basically redundant. Since we are removing multifunctional genes to test for robustness, it might seem like this should downgrade the multifunctional functions as well. Such reasoning underlies at least some enrichment software’s pre-filtering. However, as we noted earlier (**Figure 3C**), functions returned because they are multifunctional are likely to be robust, since the aggregate of any remaining biological signals will yield such functions. In contrast, the functions susceptible to removal of a single multifunctional gene are (paradoxically) not likely to be very multifunctionally biased themselves. The multifunctional gene is simply a good bet to affect any given functional enrichment, and then non-robust functions will be disrupted.

Thus far the analysis simply confirms that the robustness and uniqueness heuristics are behaving as expected, overall, on the four case studies across 17 different enrichment methods. But the true power of this assessment is to help understand the behaviour of specific methods. We divide the enrichment tools into those which apply standard statistical tests (enrichment only – 5 methods considered “non-correcting”) and those which attempt to improve on standard approaches (through statistical corrections or filtering of GO terms, etc., 12 “correcting” methods, see **Table 1**). Looking at whether two or more algorithms report the same GO terms enriched for a given study, the non-correcting algorithms overlap (at least two algorithms report) on average 58.3% (across studies, SE ~9%) of functions; in contrast, the correcting methods overlap in on average 24.2% of functions (when downsampled to the same number of algorithms; SE ~5%). And, as expected, the union of functions ever reported as enriched is much higher for the correcting algorithms (average downsampled ~58.2%) than the non-correcting (37.2%). Thus, it is hard to see any methodological convergence in advances among enrichment methods and methodological variance is likely to make results even less robust. While the methods themselves are diverse, their output can be heuristically understood in a consistent way in the light of our multifunctionality assessments. Partitioning our multifunctionality-based assessment by the enrichment method class (**Figure 5D**), we can see that ‘correcting’ methods are much more likely to return robust and unique results (average ~30.7%, SD 10.4%) in contrast to non-correcting methods (average ~16.7%, SD 6.7%), as described in **Table 2** (more details in **Supplementary Table 2** and **3**).

**Table 1.**
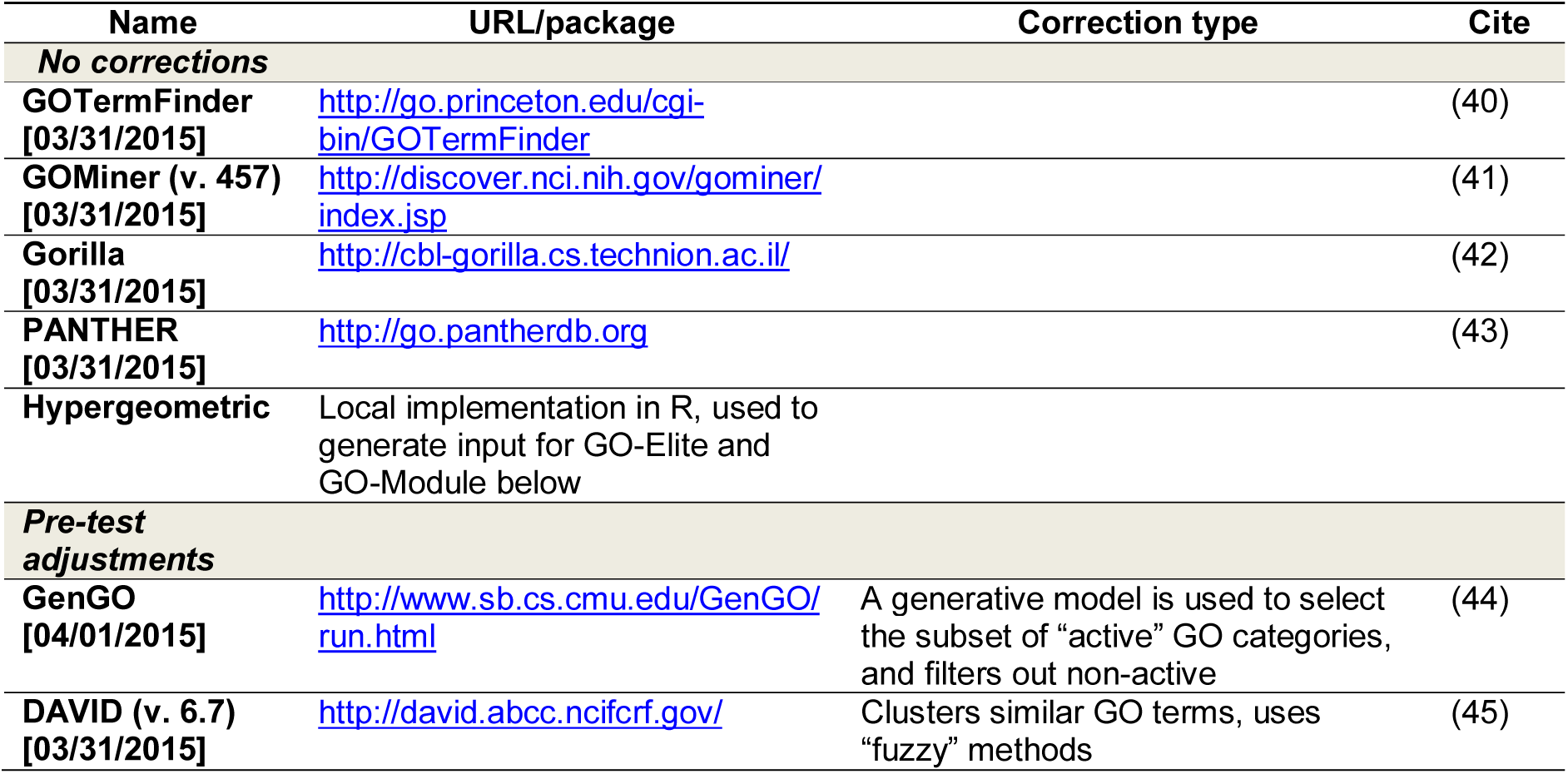
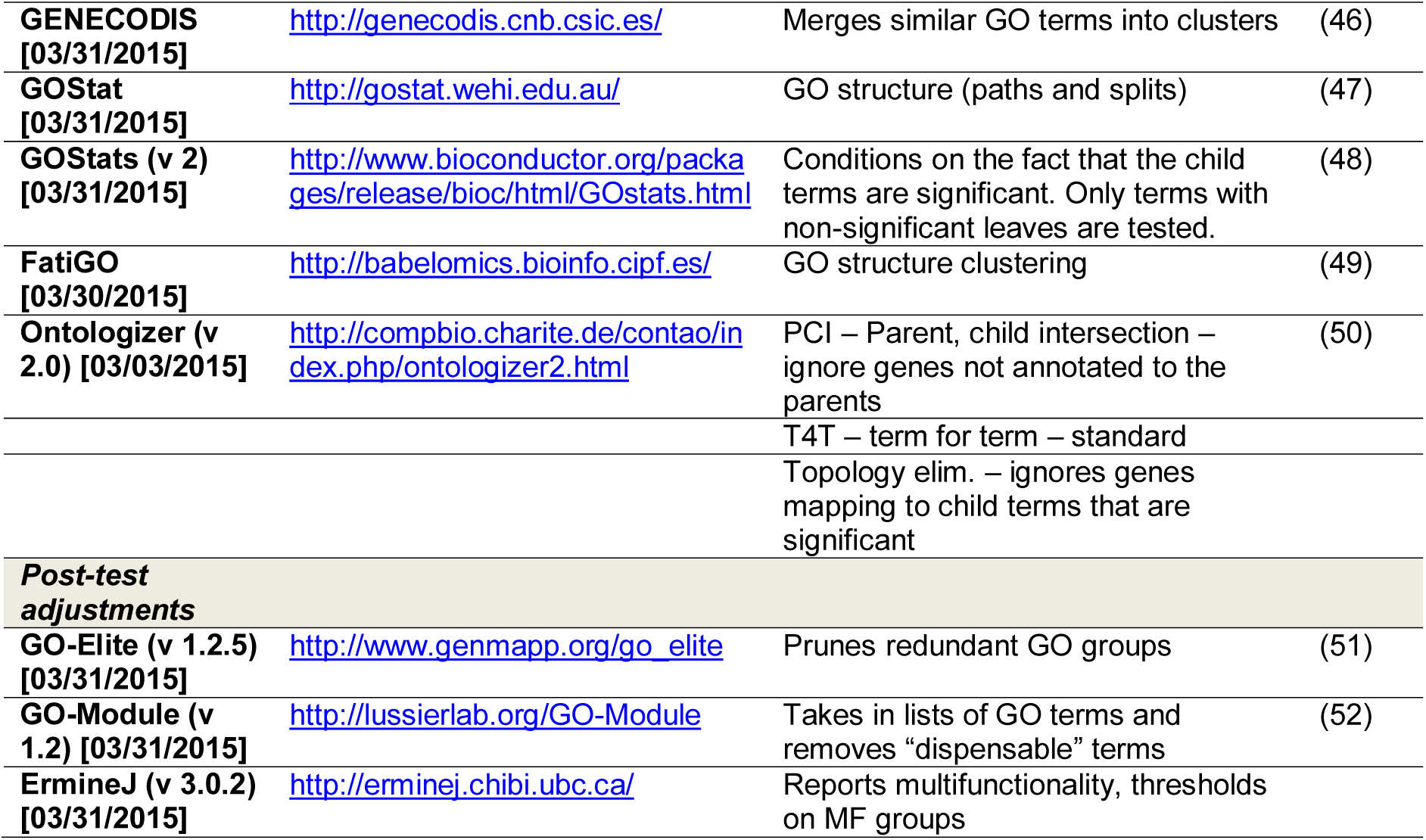
Methods used and their correction types

**Table 2.** Significant GO term result overlaps of the 17 algorithms on the four case studies

|  | Enrichments only |  | Corrected |  | Down-sampling<br>(5) | SD |
| --- | --- | --- | --- | --- | --- | --- |
|  | Average | SD | Average | SD |  |  |
| **2 or more methods** |  |  |  |  |  |  |
| All results | 58.4% | 9.2% | 52.4% | 5.3% | 24.2% | 10.7% |
| Both robust & unique | 37.4% | 17.9% | 29.0% | 8.5% | 15.9% | 8.1% |
| **Enriched across all studies (union)** |  |  |  |  |  |  |
| All results | 37.2% | 4.1% | 97.3% | 1.1% | 58.2% | 22.9% |
| Both robust & unique | 29.2% | 4.8% | 91.6% | 4.9% | 48.0% | 7.9% |
| **Enriched per study (averaged)** |  |  |  |  |  |  |
| Both robust & unique | 16.7% | 10% | 30.7% | 26% | 30.5% | 6% |

### Understanding the Gene Ontology using Multifunctionality

The correction-based algorithms we have characterized start with the GO annotations and then attempt to moderate the impacts of properties like annotation bias on a particular analysis. An alternative, as described in the introduction, would be to alter or filter the GO hierarchy to reduce multifunctionality bias and apply that “improved” GO to all analyses. The effect of such manipulations can now be evaluated in a useful way using our methods. In the following two sub-sections, we exploit this ability to perform general assessments which characterize both fine-scale features and the general architecture of GO.

#### Dissecting the Gene Ontology and its Annotations

To assess the relative contribution to multifunctionality bias of species (mouse or human), annotation codes, GO domain (e.g., biological process), and GO relation type (e.g., part_of), we built 512 alternative versions of the Gene Ontology and its annotations (collectively coined as “alt-GOs”, see methods). Returning to an observation presented earlier for the default human GO, we calculated the gene multifunctionality scores for each alternative GO, yielding a single ranked gene list. Assessing the fraction of GO terms enriched in this list at an FDR<0.05 and FDR<1E-10 gives a feel for how multifunctionality-biased the annotations are. Recall from our earlier results that in a default human GO annotation set, nearly all GO terms are enriched at FDR < 0.05 and over one third are at FDR <1E-10 (described in the above results section **Tests for multifunctional effects in gene set enrichment**). For the alt-GOs we extend this analysis to evaluate enrichment of each possible pair of alt-GOs (create the ranking with one GO, evaluate enrichment using another). This was done for all possible pairs including the simple self-comparison (**Figure 6A** for FDR<0.05 and **Supplementary Figure 8** for FDR<1E-10).

**Figure 6.**
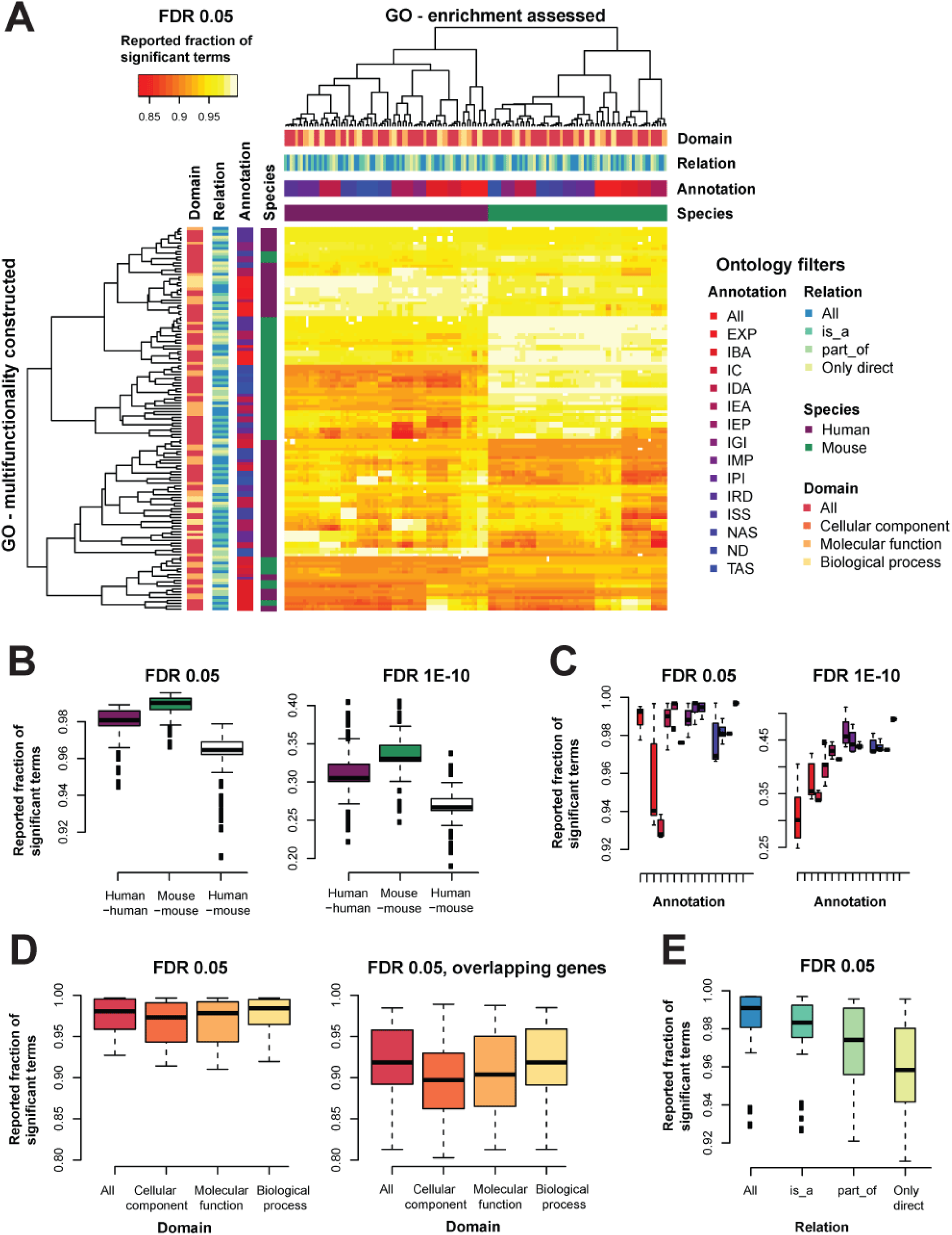
Multifunctionality bias is a robust feature of GO. Different versions of GO, filtered for different annotation properties were built and compared. (A) Heatmap of the fraction of GO terms enriched in a list of genes ordered by multifunctionality estimated from a different version of GO at an FDR of 0.05. We see a clear structure within a given ontology, with clustering by species (mouse in green, human in purple), then by annotation (shades from red to blue), then by domain (shades from orange to yellow) and some clustering by the relation (shades from blue to green). The color codes listed in the key are used consistently throughout the rest of the panels. (B) The fractions for the original human GO on all the other human GO derivatives, the original mouse GO on all the mouse derivatives, and all the human GOs on the mouse GOs. This is shown for both the FDR<0.05 and FDR<1E-10. (C) Taking each multifunctionality list and calculating the enrichment on “itself”, we see for the GO’s conditioned on annotation codes follow an upward trend to be more enriched as the annotations are more reliable. (D) For domains, this is fairly stable. (E) For relations, it is the opposite, as we become more strict, we lose the multifunctionality bias, moderately.

Across all the possible pairs of versions of GO and its annotations, the multifunctionality bias remains very high (**Figure 6A**). All the versions yield quite high reported fractions of enriched terms (>0.85; **Figure 6A**). There is also clear structure such that fractions of enriched terms within a given ontology (columns) are perfectly clustered by species first, then evidence code, with moderate clustering by domain and little by relation. Multifunctionality is thus, itself, robustly estimated from the various ontologies and so varies only modestly from GO row to row.

When split by species (**Figure 6B**), the multifunctionality from the complete GO and all evidence code set is enriched (FDR<0.05) in every derivative ontology’s annotated gene sets for greater than 94% of GO terms (median 98%). The trend in mouse is slightly stronger with all derivative ontologies being enriched in more than 96% of GO terms. We can perform the same assessment across species, using the original multifunctionality scores from each species to test for enrichment in the other; this modestly lowers reported enrichments (minimum 90%). The proportions scale similarly to our original assessment in that fractions at FDR<1E-10 show broadly similar trends centering at around 30% of terms being enriched in each derivative ontology (not shown).

If we are to consider each derivative ontology only with respect to its own multifunctionality, then our evaluation reduces to the diagonal of the heatmap in **Figure 6A**, and performance subject to evidence code choice (**Figure 6C**), domain (**Figure 6D**), and relation (**Figure 6E**) can each be evaluated. Some interesting patterns are clearly evident, with different evidence codes having different roles in extreme multifunctionalities; interestingly, traceable author statements (TAS) has relatively high multifunctionality bias while simply using all evidence codes has far less. This can be seen in the left panel of **Figure 6C,** where TAS (far right) has mean 47.4% (median 48.6%) of terms enriched at FDR<1E-10, while using all evidence codes (far left of the same panel) yields only mean 30.2% (median 28.7%) of terms enriched at the same level; this multifunctionality bias in TAS may reflect the focused annotation efforts (or biases) by curators. The more information via propagation used in the relational structure of GO, the stronger the multifunctionality bias (**Figure 6E**). However, we think the variation in values is actually quite modest given the enormous redundancy one can imagine propagation induces. This seems more a feature of the underlying biology or our knowledge of it rather than a problem with GO’s structure.

#### Rebuilding the Gene Ontology and its Annotations

Just as we can use the multifunctionality heuristic to readily parse GO and its annotations in fine detail, we can consider how even more vastly altered Gene Ontologies would behave through this heuristic. In the following section, we consider four radically different versions of the Gene Ontology and annotations (summarized in **Figure 7**).

**Figure 7.**
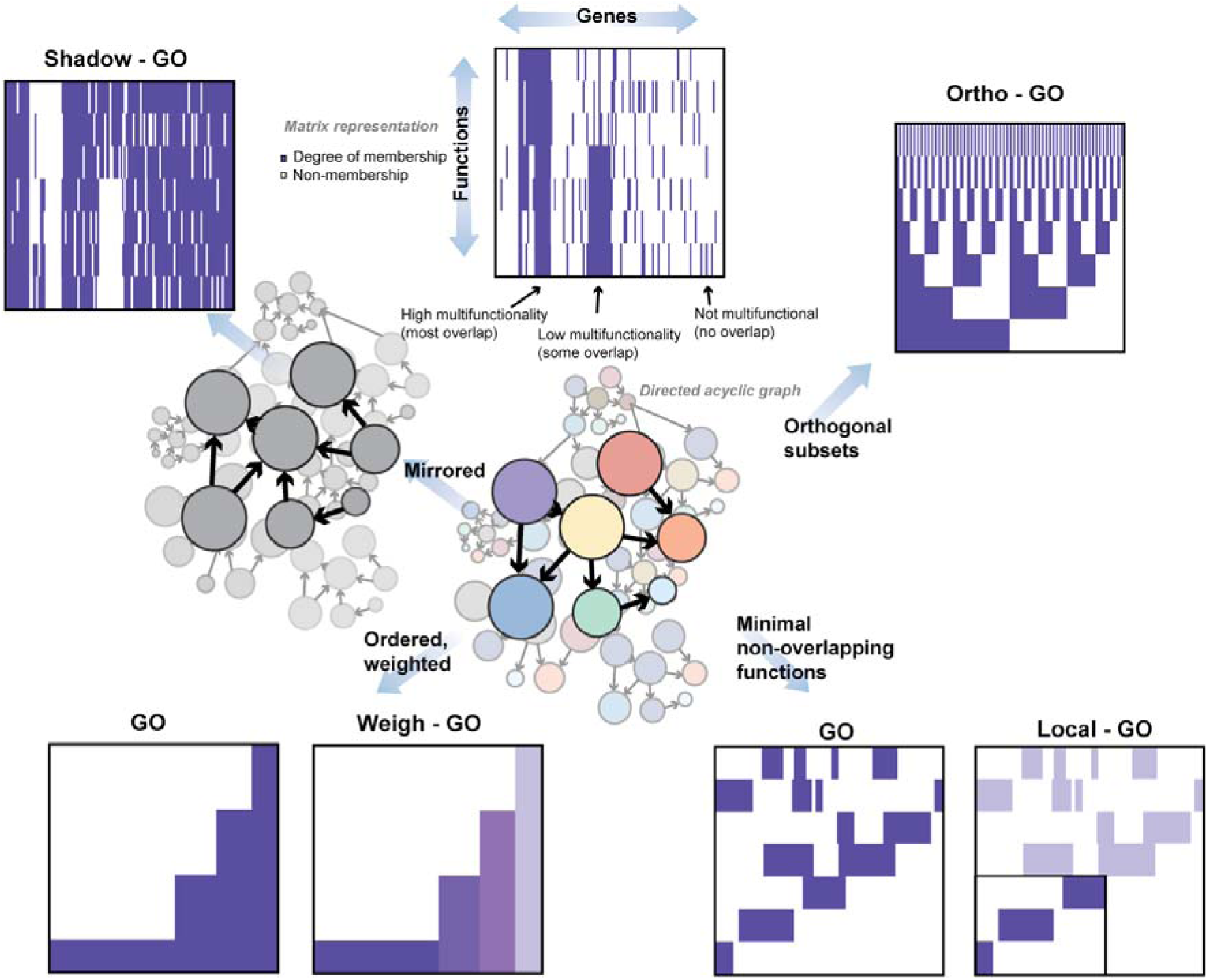
Alternate hypothetical GOs to assess multifunctionality. Constraining the ontology to reduce or change multifunctionality can be done by “Shadowing” GO to obtain members “not” in GO. Selecting orthogonal groups to force minimal overlaps (Ortho-GO), or weighting the genes (Weigh-GO) so that overall membership is the same also constrains multifunctionality. A targeted version (Local-GO) has a more specific and hypothesis driven basis, where a known function is pre-selected, and other GO groups are selected that do not intersect with each other.

In our first version of GO, every Gene Ontology term brings into existence its opposite to which genes would additionally be annotated as “not” being members. This yields a logically consistent GO, and forces all genes to have the same number of annotations (one for each GO term, as to whether it belongs or not). The main other effect is to formalize a closed world assumption on GO, which is consistent with how GO is normally applied. Despite having no annotation bias, multifunctionality scores from the Shadow-GO are still highly correlated with those derived from the original GO (Spearman’s rank correlation r_s_ ~ 0.69). Shadow-GO also suffers from almost the same enrichment problems as default GO with almost 97% of gene sets enriched at FDR<0.05 and 37% at FDR<1E-10. However, unlike in the original assessment, where sidedness had no impact, the multifunctionality ranking is now highly sided, with only 50% of GO terms being positively enriched. This is almost fully accounted for by 99% of the original GO terms being positively enriched with only a few of the new shadow terms being positively enriched. This is striking since the calculation of multifunctionality is “unaware” of the structure of GO and simply sees the annotated sets. Conceptually the modest impact the GO shadowing has makes sense: Shadow-GO is at least as redundant as the original GO and is only concealing its bias from a trivial assessment. We consider two additional hypothetical-GOs, Ortho-GO and Weigh-GO (see supplement: “Rebuilding the Gene Ontology…”), both of which may be thought of as elaborations of Shadow-GO, in one case by constructing gene sets to be orthogonal and in the other by weighting gene membership within existing gene sets. In both cases, while annotation bias is largely gone, the utility of multifunctionality in understanding “expected” enrichment results is still very high.

Our final version of GO, “Local-GO”, also discards GO as a universal tool and asks only if we can construct local non-overlapping sets that “work”. This is close to the premise of GO-slim and one might imagine it as pre-registering a function of interest for a given experiment and then having that function define which other sets are independent enough to also be tested. One obvious limitation of this approach is that not all of the genes originally possessing some function will now have one annotated. Indeed we see this: the 200-local-GO attached annotations to only 14% of originally covered genes and the 1000-local-GO attached annotations to only 45% of originally annotated genes (see methods). While 68% of GO groups were significantly enriched by multifunctionality on 200-local-GO (and 52% on 1000-local-GO), almost none were very significantly enriched (~1% 200-local-GO, ~2% 1000-local-GO FDR<1E-10), even with the more modest multiple hypothesis test correction. Thus, using pre-registration, semantic filtering and extreme enrichment thresholds would seem to be a potential improvement in ensuring results were biologically specific.

### Implementation of multifunctionality reporting in ErmineJ

The approaches we describe are general enough that they can be adapted to any gene set analysis method. To permit biologists to rapidly benefit, we have integrated new features in ErmineJ (version 3.0) that expose information on multifunctionality to users. ErmineJ (22,23) is open source desktop software implemented in the Java programming language that affords a point-and-click interface for enrichment analysis with extensive visualization features, as well as programmatic and scriptable interfaces. Our philosophy in designing the multifunctionality features of ErmineJ 3.0 is to make it clear which results are sensitive to multifunctionality, rather than to focus on corrected results as such. Users can then decide to filter or re-rank the results based on multifunctionality effects. **Figure 8** is a screenshot of the ErmineJ 3.0 interface illustrating the presentation of multifunctionality effects for the hypoxia case study. ErmineJ also provides diagnostic plots of multifunctionality which can be useful for detecting how biased the user’s data is before analysis. The new ErmineJ features are documented at http://erminej.chibi.ubc.ca/.

**Figure 8.**
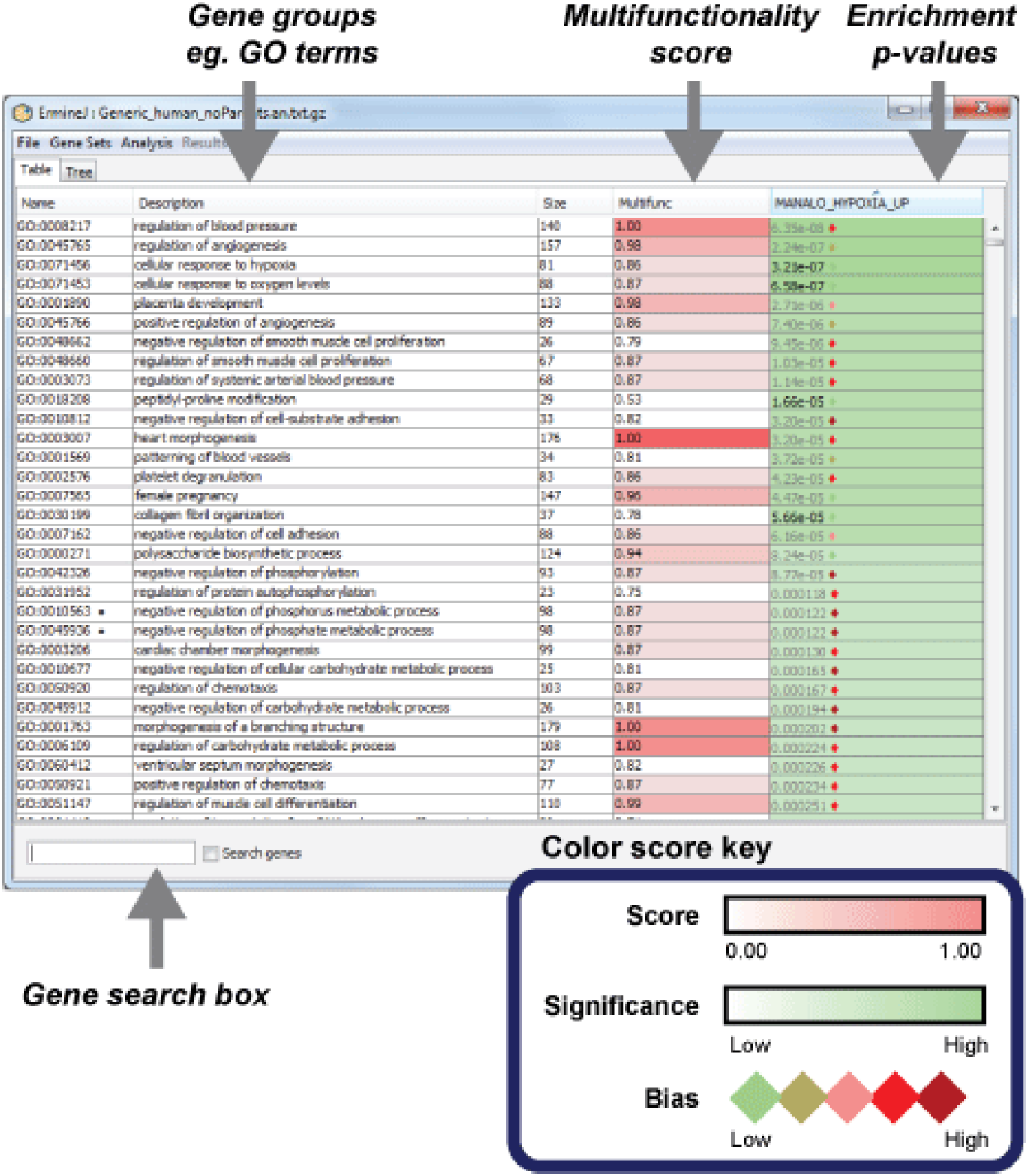
Screen shot of the ErmineJ interface showing multifunctionality-related features. The results of the enrichment analysis for the Manalo case study are shown. Each row in the table is a gene group. The right-most column shows the enrichment results as p-values, with different tints of green indicating strength of enrichment. The degree to which the result is sensitive to multifunctionality correction is indicated by a diamond next to the p-value, with red indicating the highest sensitivity. The p-value is shown in grey if it would not be significant at an FDR of 0.1 after multifunctionality correction. The second column from the left shows the multifunctionality of the GO term, which darker tints of red indicating stronger bias. Note that the most multifunctional GO terms are not necessarily the ones which have the strongest effect of correction.

## DISCUSSION

Gene set analysis allows us to make statements ascribing an experimental result to changes in underlying gene-based functions. This is of tremendous scientific value if it works and can be correspondingly damaging if it does not. No consensus has emerged as to the correct way to perform function enrichment, and this is impossible to resolve without accepted gold standards, which are currently not attainable. Rather than tackle this issue, we have suggested a generally applicable heuristic test to assess if function enrichment is working reasonably in terms of robustness and uniqueness. We showed that both of these properties are highly associated with gene multifunctionality (as operationalized by gene sets). In the simplest version of our approach, when testing for enrichment of a gene set, one simply removes the most multifunctional gene from the data and reruns the analysis in whatever software tool being used. This can be repeated until either the notable results vanish or the experimenter is comfortable that the results are robust and can be reported as such, e.g., “Our 350 candidate genes were enriched for synaptic activity, a property robust to the removal of the 5 most multifunctional genes”. By considering robustness over multifunctional genes, weak signals can still be considered significant if they are unusual (of low prior probability).

In contrast to our approach, attempts to improve enrichment methods to better recover the true functions by fixing the underlying enrichment tests or determining the underlying dependencies of functions are trying to address ill-posed problem given the current incompleteness of annotation data. We do not know how functions combine (linearly?), whether an absence of annotations reflects an annotation of absence (a necessary and untrue assumption), or indeed, the true null distribution for many experimental designs. We propose that the best we can hope is to assess the conditions under which enrichment results are not sensitive to these likely confounds. An important result from our analysis is that the methods that do attempt to correct for biases and lack of gene independence have succeeded even while ending up with highly variable output. We think this is a generally important principle that may be fruitfully generalize to other areas of bioinformatics research: Even treating the methods as black boxes, uniqueness and robustness are fundamental properties which can serve to benchmark methods. Our formal assessment of this for enrichment focusses on multifunctionality, but these ideas will find alternate expression in different areas of research and serve as an important alternative to developing methods based on assessment in specific gold standards, which can result in fieldwide overfitting (39). Our assessment of the current state of corrections in enrichment – that they work, modestly – was both encouraging and surprising to us. Combined with our evaluation of hypothetical GOs, we feel these demonstrations provide strong evidence of the intuitive significance of our approaches and create a strong argument for making multifunctionality considerations a routine aspect of enrichment analysis.

We have shown that gene multifunctionality has a major impact on the biological interpretability of functional enrichment analysis, and presented algorithms that improve the interpretability of results, often dramatically. Enrichment analysis is already extremely widely used, and we suggest that accounting for multifunctionality will make it a more attractive approach for interpreting genomics studies. The availability of the implementations in ErmineJ 3.0 put these approaches in the hands of researchers. The algorithms are also simple to implement and general, and could easily be adopted for use in other software packages.

## REFERENCES

1. Tavazoie, S., Hughes, J.D., Campbell, M.J., Cho, R.J. and Church, G.M. (1999) Systematic determination of genetic network architecture. Nat. Genet., 22, 281–285.

2. Pavlidis, P., Lewis, D.P. and Noble, W.S. (2002) Exploring gene expression data with class scores. Pac Symp Biocomput, 474–485.

3. Mootha, V.K., Lindgren, C.M., Eriksson, K.F., Subramanian, A., Sihag, S., Lehar, J., Puigserver, P., Carlsson, E., Ridderstrale, M., Laurila, E. et al. (2003) PGC-1alpha-responsive genes involved in oxidative phosphorylation are coordinately downregulated in human diabetes. Nat. Genet., 34, 267–273.

4. Khatri, P., Draghici, S., Ostermeier, G.C. and Krawetz, S.A. (2002) Profiling gene expression using onto-express. Genomics, 79, 266–270.

5. Ashburner, M., Ball, C.A., Blake, J.A., Botstein, D., Butler, H., Cherry, J.M., Davis, A.P., Dolinski, K., Dwight, S.S., Eppig, J.T. et al. (2000) Gene ontology: tool for the unification of biology. The Gene Ontology Consortium. Nat. Genet., 25, 25–29.

6. Kanehisa, M. and Goto, S. (2000) KEGG: kyoto encyclopedia of genes and genomes. Nucleic Acids Res., 28, 27–30.

7. Hamosh, A., Scott, A.F., Amberger, J.S., Bocchini, C.A. and McKusick, V.A. (2005) Online Mendelian Inheritance in Man (OMIM), a knowledgebase of human genes and genetic disorders. Nucleic Acids Res., 33, D514–D517.

8. Gillis, J. and Pavlidis, P. (2011) The impact of multifunctional genes on “guilt by association” analysis. PloS one, 6, e17258.

9. Alterovitz, G., Xiang, M., Mohan, M. and Ramoni, M.F. (2007) GO PaD: the Gene Ontology Partition Database. Nucleic Acids Res., 35, D322–327.

10. Zambon, A.C., Gaj, S., Ho, I., Hanspers, K., Vranizan, K., Evelo, C.T., Conklin, B.R., Pico, A.R. and Salomonis, N. (2012) GO-Elite: a flexible solution for pathway and ontology over-representation. Bioinformatics, 28, 2209–2210.

11. Davis, M.J., Sehgal, M.S. and Ragan, M.A. (2010) Automatic, context-specific generation of Gene Ontology slims. BMC Bioinformatics, 11, 498.

12. Geifman, N., Monsonego, A. and Rubin, E. (2010) The Neural/Immune Gene Ontology: clipping the Gene Ontology for neurological and immunological systems. BMC Bioinformatics, 11, 458.

13. Jin, B. and Lu, X. (2010) Identifying informative subsets of the Gene Ontology with information bottleneck methods. Bioinformatics, 26, 2445–2451.

14. Yang, X., Li, J., Lee, Y. and Lussier, Y.A. (2011) GO-Module: functional synthesis and improved interpretation of Gene Ontology patterns. Bioinformatics, 27, 1444-1446.

15. Jantzen, S.G., Sutherland, B.J., Minkley, D.R. and Koop, B.F. (2011) GO Trimming: Systematically reducing redundancy in large Gene Ontology datasets. BMC Res. Notes., 4, 267.

16. Supek, F., Bosnjak, M., Skunca, N. and Smuc, T. (2011) REVIGO summarizes and visualizes long lists of gene ontology terms. PloS one, 6, e21800.

17. Lu, Y., Rosenfeld, R., Simon, I., Nau, G.J. and Bar-Joseph, Z. (2008) A probabilistic generative model for GO enrichment analysis. Nucleic Acids Res., 36, e109.

18. Tarca, A.L., Draghici, S., Bhatti, G. and Romero, R. (2012) Down-weighting overlapping genes improves gene set analysis. BMC Bioinformatics, 13, 136.

19. Glass, K. and Girvan, M. (2014) Annotation Enrichment Analysis: An Alternative Method for Evaluating the Functional Properties of Gene Sets. Sci. Rep., 4, 4191.

20. Khatri, P., Bhavsar, P., Bawa, G. and Draghici, S. (2004) Onto-Tools: an ensemble of web-accessible, ontology-based tools for the functional design and interpretation of high-throughput gene expression experiments. Nucleic Acids Res., 32, W449–456.

21. Subramanian, A., Tamayo, P., Mootha, V.K., Mukherjee, S., Ebert, B.L., Gillette, M.A., Paulovich, A., Pomeroy, S.L., Golub, T.R., Lander, E.S. et al. (2005) Gene set enrichment analysis: a knowledge-based approach for interpreting genome-wide expression profiles. Proc Natl Acad Sci U S A, 102, 15545–15550.

22. Gillis, J., Mistry, M. and Pavlidis, P. (2010) Gene function analysis in complex data sets using ErmineJ. Nat. Protoc., 5, 1148–1159.

23. Lee, H.K., Braynen, W., Keshav, K. and Pavlidis, P. (2005) ErmineJ: tool for functional analysis of gene expression data sets. BMC Bioinformatics, 6, 269.

24. Meyer, L.R., Zweig, A.S., Hinrichs, A.S., Karolchik, D., Kuhn, R.M., Wong, M., Sloan, C.A., Rosenbloom, K.R., Roe, G., Rhead, B. et al. (2013) The UCSC Genome Browser database: extensions and updates 2013. Nucleic Acids Res., 41, D64–69.

25. Maglott, D., Ostell, J., Pruitt, K.D. and Tatusova, T. (2011) Entrez Gene: gene-centered information at NCBI. Nucleic Acids Res., 39, D52–57.

26. Liberzon, A., Subramanian, A., Pinchback, R., Thorvaldsdottir, H., Tamayo, P. and Mesirov, J.P. (2011) Molecular signatures database (MSigDB) 3.0. Bioinformatics, 27, 1739–1740.

27. Breslin, T., Eden, P. and Krogh, M. (2004) Comparing functional annotation analyses with Catmap. BMC Bioinformatics, 5, 193.

28. Benjamini, Y. and Hochberg, Y. (1995) Controlling the False Discovery Rate: a Practical and Powerful Approach to Multiple Testing. Journal of the Royal Statistical Society B, 57, 12.

29. Portales-Casamar, E., Ch’ng, C., Lui, F., St-Georges, N., Zoubarev, A., Lai, A.Y., Lee, M., Kwok, C., Kwok, W., Tseng, L. et al. (2013) Neurocarta: aggregating and sharing disease-gene relations for the neurosciences. BMC Genomics, 14, 129.

30. Jonquet, C., Shah, N.H. and Musen, M.A. (2009) The open biomedical annotator. Summit on translational bioinformatics, 2009, 56–60.

31. Dimmer, E.C., Huntley, R.P., Alam-Faruque, Y., Sawford, T., O’Donovan, C., Martin, M.J., Bely, B., Browne, P., Chan, W.M., Eberhardt, R. et al. (2012) The UniProt-GO Annotation database in 2011. Nucleic Acids Res., 40, D565–D570.

32. Goeman, J.J. and Buhlmann, P. (2007) Analyzing gene expression data in terms of gene sets: methodological issues. Bioinformatics, 23, 980–987.

33. Gilman, S.R., Iossifov, I., Levy, D., Ronemus, M., Wigler, M. and Vitkup, D. (2011) Rare de novo variants associated with autism implicate a large functional network of genes involved in formation and function of synapses. Neuron, 70, 898–907.

34. Levy, D., Ronemus, M., Yamrom, B., Lee, Y.H., Leotta, A., Kendall, J., Marks, S., Lakshmi, B., Pai, D., Ye, K. et al. (2011) Rare de novo and transmitted copy-number variation in autistic spectrum disorders. Neuron, 70, 886–897.

35. Schmidt-Kastner, R., van Os, J., Esquivel, G., Steinbusch, H.W. and Rutten, B.P. (2012) An environmental analysis of genes associated with schizophrenia: hypoxia and vascular factors as interacting elements in the neurodevelopmental model. Mol. Psychiatry, 17, 1194–1205.

36. Manalo, D.J., Rowan, A., Lavoie, T., Natarajan, L., Kelly, B.D., Ye, S.Q., Garcia, J.G. and Semenza, G.L. (2005) Transcriptional regulation of vascular endothelial cell responses to hypoxia by HIF-1. Blood, 105, 659–669.

37. Pardo, M., Lang, B., Yu, L., Prosser, H., Bradley, A., Babu, M.M. and Choudhary, J. (2010) An expanded Oct4 interaction network: implications for stem cell biology, development, and disease. Cell Stem Cell, 6, 382–395.

38. Gillis, J. and Pavlidis, P. (2012) “Guilt by association” is the exception rather than the rule in gene networks. PLoS Comput. Biol., 8, e1002444.

39. Verleyen, W., Ballouz, S. and Gillis, J. (2015) Positive and negative forms of replicability in gene network analysis. Bioinformatics.

40. Boyle, E.I., Weng, S., Gollub, J., Jin, H., Botstein, D., Cherry, J.M. and Sherlock, G. (2004) GO:: TermFinder—open source software for accessing Gene Ontology information and finding significantly enriched Gene Ontology terms associated with a list of genes. Bioinformatics, 20, 3710–3715.

41. Zeeberg, B.R., Feng, W., Wang, G., Wang, M.D., Fojo, A.T., Sunshine, M., Narasimhan, S., Kane, D.W., Reinhold, W.C. and Lababidi, S. (2003) GoMiner: a resource for biological interpretation of genomic and proteomic data. Genome Biology, 4, R28.

42. Eden, E., Navon, R., Steinfeld, I., Lipson, D. and Yakhini, Z. (2009) GOrilla: a tool for discovery and visualization of enriched GO terms in ranked gene lists. BMC Bioinformatics, 10, 48.

43. Mi, H., Muruganujan, A., Casagrande, J.T. and Thomas, P.D. (2013) Large-scale gene function analysis with the PANTHER classification system. Nat. Protoc., 8, 1551–1566.

44. Lu, Y., Rosenfeld, R., Simon, I., Nau, G.J. and Bar-Joseph, Z. (2008) A probabilistic generative model for GO enrichment analysis. Nucleic Acids Research, 36, e109.

45. Huang, D.W., Sherman, B.T. and Lempicki, R.A. (2008) Systematic and integrative analysis of large gene lists using DAVID bioinformatics resources. Nat. Protoc., 4, 44–57.

46. Carmona-Saez, P., Chagoyen, M., Tirado, F., Carazo, J.M. and Pascual-Montano, A. (2007) GENECODIS: a web-based tool for finding significant concurrent annotations in gene lists. Genome Biology, 8, R3.

47. Beißbarth, T. and Speed, T.P. (2004) GOstat: find statistically overrepresented Gene Ontologies within a group of genes. Bioinformatics, 20, 1464–1465.

48. Falcon, S. and Gentleman, R. (2007) Using GOstats to test gene lists for GO term association. Bioinformatics, 23, 257–258.

49. Al-Shahrour, F., Díaz-Uriarte, R. and Dopazo, J. (2004) FatiGO: a web tool for finding significant associations of Gene Ontology terms with groups of genes. Bioinformatics, 20, 578–580.

50. Bauer, S., Grossmann, S., Vingron, M. and Robinson, P.N. (2008) Ontologizer 2.0—a multifunctional tool for GO term enrichment analysis and data exploration. Bioinformatics, 24, 1650–1651.

51. Zambon, A.C., Gaj, S., Ho, I., Hanspers, K., Vranizan, K., Evelo, C.T., Conklin, B.R., Pico, A.R. and Salomonis, N. (2012) GO-Elite: a flexible solution for pathway and ontology over-representation. Bioinformatics, 28, 2209–2210.

52. Yang, X., Li, J., Lee, Y. and Lussier, Y.A. (2011) GO-Module: functional synthesis and improved interpretation of Gene Ontology patterns. Bioinformatics, 27, 1444–1446.

